# A zebrafish model of nicotinamide adenine dinucleotide (NAD^+^) deficiency-derived congenital disorders

**DOI:** 10.1101/2025.01.10.632366

**Authors:** Visakuo Tsurho, Carla Gilliland, Jessica Ensing, Elizabeth Vansickle, Nathan J. Lanning, Paul R. Mark, Stephanie Grainger

## Abstract

Congenital NAD deficiency disorder (CNDD) is a multisystem condition in which cardiac, renal, vertebral, and limb anomalies are most common, but anomalies in all organ systems have been identified. Patients with this condition have biallelic pathogenic variants involving genes in the nicotinamide adenine dinucleotide (NAD^+^) synthesis pathway leading to decreased systemic NAD^+^ levels. CNDD anomalies mimic the clinical features described in vertebral-anal-cardiac-tracheoesophageal fistula-renal-limb (VACTERL) association raising the possibility that CNDD and VACTERL association possess similar underlying causes. However, the mechanism by which NAD^+^ deficiency causes CNDD developmental anomalies has not been determined, nor has NAD^+^ deficiency been definitively linked to VACTERL association. Therefore, additional animal models amenable to detailed observation of embryonic development are needed to address the causes and progression of congenital anomalies in both CNDD and VACTERL association. Here, we describe a zebrafish model of NAD^+^ disruption to begin to model CNDD and VACTERL association phenotypes, assessing developmental anomalies in real-time. Treatment of zebrafish embryos with 2-amino-1,3,4-thiadiazole (ATDA), a teratogen known to disrupt NAD^+^ metabolism, resulted in neural tube, craniofacial, cardiac, and tail defects. These defects were rescued by the administration of nicotinamide (NAM) in a dose-dependent manner. Our work establishes zebrafish as a useful model for investigating the mechanistic causes and developmental dynamics of CNDD and VACTERL association. Further, as VACTERL association has been linked to teratogens, our zebrafish model provides a platform to assess these agents.

**One sentence summary:** NAD(H) deficiency causes multiple congenital anomalies in zebrafish.

## Introduction

NAD^+^, its reduced form (NADH), and its phosphorylated forms (NADP^+^/NADPH) serve as coenzymes, cosubstrates, and cofactors for numerous critical biochemical reactions (Migaud et al., 2024). Therefore, NAD^+^ levels are tightly regulated both intracellularly and systemically, and multiple cellular mechanisms exist for maintaining oxidized and reduced ratios of NAD+ and their phosphorylated forms (Migaud et al., 2024; Xie et al., 2020). It is difficult to study the impacts of NAD^+^ loss genetically because it is synthesized *de novo* through the kynurenine and Preiss-Handler pathways, as well as through a salvage pathway (Fig. 1). In addition to this, multiple dietary precursors, including tryptophan, nicotinic acid, nicotinamide (NAM), and nicotinamide riboside feed NAD^+^ biosynthesis (Xie et al., 2020) (Fig. 1). Complete ablation of NAD^+^ synthesis from all pathways is likely embryonic lethal, and partial loss of NAD+ in mice is incompletely penetrant (Bozon et al., 2024; Cuny et al., 2020; Shi et al., 2017). Because of this, it has been difficult to interpret the requirements for NAD^+^ in animal development.

**Figure 1.**
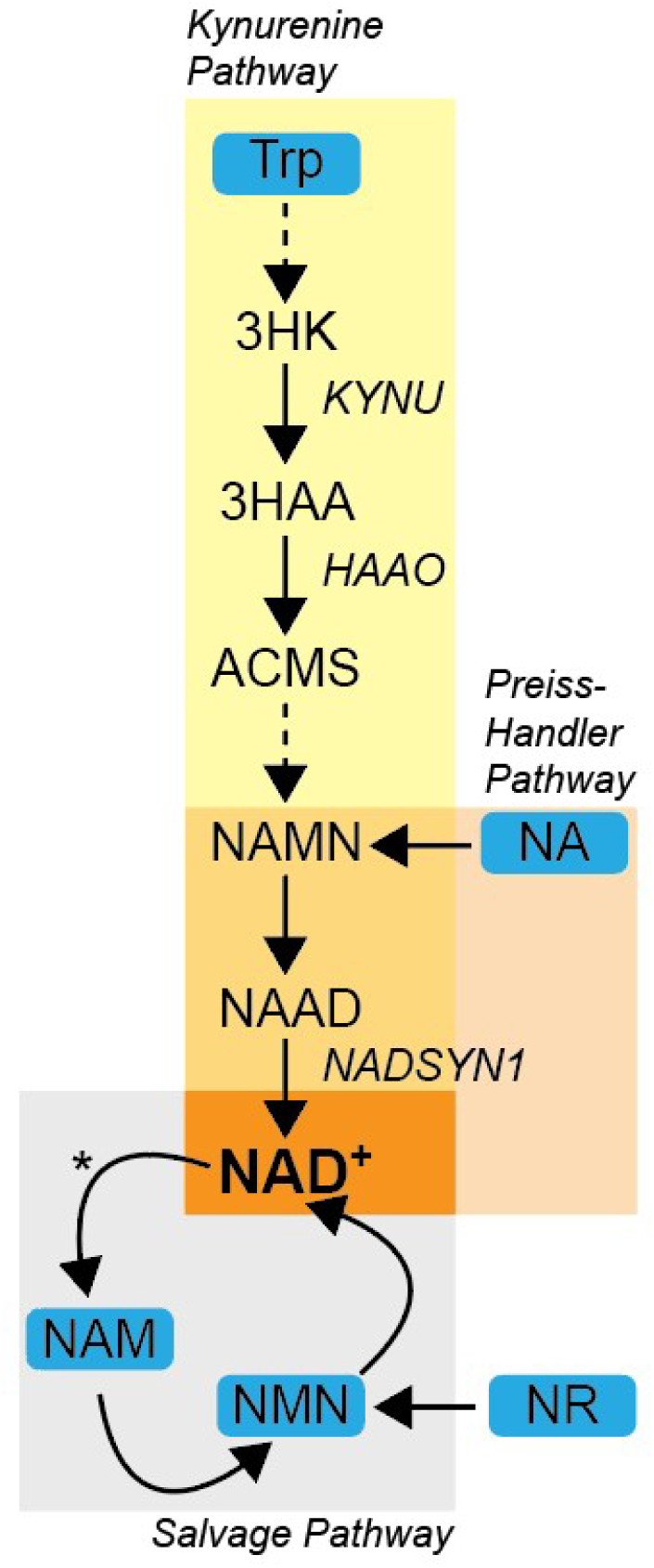
NAD^+^ biosynthesis and salvage. Organismal NAD^+^ is derived from dietary precursor sources indicated with blue rectangle backgrounds. NAD^+^ precursors flux through the kynurenine (yellow) and Preiss-Handler (orange) biosynthetic pathways or are incorporated into the salvage pathway (gray). The bulk of cellular NAD^+^ is derived from the salvage pathway. NAD^+^ is consumed as a substrate (asterisk in the salvage pathway) by enzymes such as PARPs and sirtuins. Loss-of-function mutations to *KYNU, HAAO*, and *NADSYN1* genes, which encode enzymes in the biosynthetic pathways, result in NAD^+^ depletion and CNDD.

Human congenital defects associated with NAD^+^ deficiency demonstrate the requirement for NAD^+^ in proper developmental programs. Deficient NAD^+^ synthesis or dysregulated metabolism lead to skin, intellect, hearing, cardiac, renal, skeletal, or retinal disorders (Zapata-Pérez et al., 2021). Congenital NAD deficiency disorder (CNDD) is a multisystem condition caused by biallelic variants in the NAD^+^ Synthesis Pathway genes kynurinase (KYNU), 3-hydroxyanthranilate 3,4-dioxygenase (HAAO), or NAD synthetase 1 (NADSYN1) (P. Mark & Dunwoodie, 2023). The most common features of this condition are cardiac, renal, vertebral, and limb anomalies, mimicking the clinical features described in VACTERL association, in addition to short stature and developmental delay. Anomalies in other organ systems have been identified, including neurologic, respiratory, gastrointestinal, sensory, and endocrine, therefore impacting the development of derivatives from all primary germ layers (P. R. Mark, 2022). There are a number of recurrent multiple malformation conditions which overlap in phenotype with VACTERL association and CNDD, including limb-body wall complex, pentalogy of Cantrell, omphalocele-exstrophy-imperforate anus-spinal defects complex, oculoauriculovertebral spectrum, Mullerian duct aplasia-renal anomalies-cervicothoracic somite dysplasia, sirenomelia, and urorectal septum malformation sequence (Adam et al., 2020; P. R. Mark, 2022). Although a genetic cause for these has not yet been identified, it has been hypothesized that low levels of NAD^+^ during development could cause these conditions (P. R. Mark, 2022; Shi et al., 2017; Vander Heiden, 2017). However, this has been difficult to show using animal modeling; this is likely at least in part due to the incomplete penetrance of genetic mutants in the absence of maternal dietary restriction in mouse models (Bozon et al., 2024; Cuny et al., 2020; Shi et al., 2017). Taken together, these studies indicate that loss of NAD^+^ during development can lead to diverse congenital abnormalities across germ layers.

We have now developed a zebrafish model using ATDA to induce cardiac, craniofacial, tail, body length, and neural tube defects, all reminiscent of CNDD and VACTERL. These defects were rescued by NAM administration in a dose-dependent fashion, supporting a requirement for NAD^+^ in proper development. Our work establishes zebrafish as a useful model for understanding the ontogeny of CNDD and VACTERL association anomalies and other developmental complications arising from NAD^+^ deficiency.

## Materials and Methods

### Zebrafish husbandry, spawning, and drug treatment

Zebrafish were maintained and propagated according to Van Andel Institute and local Institutional Animal Care and Use Committee policies. AB* zebrafish were used in all experiments. Wild-type adult zebrafish (*Danio rerio*) 3–8 months old were reared and maintained at a temperature of 28 ± 0.5 °C and a photoperiod of 14 h:10 h light:dark cycle. Continuous water recirculation with conductivity of 700 ± 100 μS/cm, pH at 7.4 ± 0.5, and dissolved oxygen ≥ 95% saturation were maintained.

Embryos were spawned and collected using standard methods and grown at 28.5° C. The clutches of embryos were rinsed to remove debris and redistributed into 10 cm plates with 50 embryos per treatment group, in 30 mL of Essential 3 (E3) medium (5 mM NaCl, 0.17 mM KCl, 0.33 mM CaCl2, 0.33 mM MgSO4, 10^−5^% Methylene Blue). Unfertilized eggs, embryos with cleavage anomalies, and embryos with injuries or other types of deformities were discarded (Rajesh and Divya, 2023). Stock solutions for 50 mg/ml ATDA (Thermo Fisher Scientific Inc., Waltham, MA) or 200 mg/ml NAM (Thermo Fisher Scientific Inc., Waltham, MA) were prepared in purified water and added to E3 culture medium as indicated in figures at 4 hours post fertilization (hpf). Embryos were examined for developmental defects under a Leica stereo microscope throughout development and assessed as described below.

### NAD(H) quantitation

Dechorionated 48 hpf embryos were rinsed three times in ice-cold 1X phosphate buffered saline (1.54 mM KH2PO4, 155 mM NaCl, 2.7 mM Na2HPO4-7H2O, pH 7.4). Following removal of all liquid, individual embryos were snap-frozen in liquid nitrogen. Each frozen embryo was then homogenized in 100 µl of NAD/NADH-Glo detection reagent (Promega, Madison, WI) multiplexed with two µl of CyQUANT (Thermo Fisher Scientific Inc., Waltham, MA) prepared according to the manufacturers’ specifications. Each homogenized sample was loaded into a 96-well black-walled, clear-bottom assay plate and luminescence and fluorescence was detected as previously described (Lanning et al., 2014). Individual NAD(H) luminescence values were normalized to condition-specific CyQUANT average values.

### Hatching and Mortality rate assessment

Hatching rate was evaluated beginning at 48 hpf and was determined by dividing the number of hatched embryos by the total number of live incubated embryos in each condition (Powers et al., 2010). The mortality rate was calculated by dividing the number of dead embryos by the total number of exposed embryos (Powers et al., 2010).

### Phenotypic outcomes and severity of developmental abnormalities

The proportion of growing embryos having developmental abnormalities was calculated for each experimental group. The severity of pericardial edema and tail malformation in developing embryos was graded individually at 48 hpf and 72 hpf using categories normal, mild, moderate, and severe as described in Table 1. Craniofacial defects were assigned to fish if they displayed greyness of the eyes, protruding forehead, no eyes, miniature skull, or facial/ocular edema. After examination, zebrafish were fixed with 4% PFA and then stored in methanol at -20° C.

**Table 1:** Pericardial Edema and Tail Malformation Grading.

|  | **Pericardial Edema** | **Tail Malformation** |
| --- | --- | --- |
| **Normal** | No Edema | No malformation |
| **Mild** | Mild edema causing the heart to contribute 1/2-1/3 of total heart sac volume. | One minor tail kink/lordosis/scoliosis; semi-oval shape. |
| **Moderate** | Moderate edema causing the heart to contribute to 1/4-1/6 of the total heart sac volume. | Two or more minor tail kinks/lordosis/scoliosis or one sharp tail kink/lordosis/scoliosis; semicircle shape. |
| **Severe** | Severe edema causing the heart to contribute to 1/8 or less of the total heart sac volume. | Kinks causing a >45° angle in the tail; supercoiled tail shape. |

### Histology

Tissues were processed using a Tissue Tek VIP 5 utilizing routine histology dehydration and paraffin infiltration practices. Tissues were embedded utilizing the Epredia Histostar embedding center. Sections were cut using Leica rotary microtomes at 5 µm thick and mounted to Epredia Colormark Plus slides. H&E staining was performed utilizing Sakura’s Tissue Tek Prisma Plus Autostainer with Tissue Tek Prisma H&E Stain Kit #1. Digital scanning was performed using the Aperio AT2 system.

### Statistics and Image Analysis

All statistics and image analysis were performed using GraphPad Prism or ImageJ.

## Results

### ATDA impairs zebrafish embryo viability and hatching rate

ATDA treatment of rats induced phenotypes resembling those of human CNDD and murine models of NAD(H) deficiency (Beaudoin, 1974, 1976; Scott et al., 1973) (Table 2) and was rescued by NAD^+^ metabolic precursors, indicating that NAD^+^ deficiency underscores these phenotypes. However, it is difficult to assess the ontogeny and progression of CNDD in these models since they are complicated by the maternal supply of NAD^+^ (Bozon et al., 2024; Cuny et al., 2023; Shi et al., 2017). Zebrafish exhibit rapid, external development and are optically transparent; therefore, developmental defects can be observed in real time and *in situ*. CNDD and VACTERL association present with multiple and varied phenotypes, perhaps related to the timing of genetic or environmental insults, in addition to the tissues impacted. Therefore, the ability dynamically assess development of these diseases is critical to developing suitable treatments and/or prevention options.

**Table 2:**
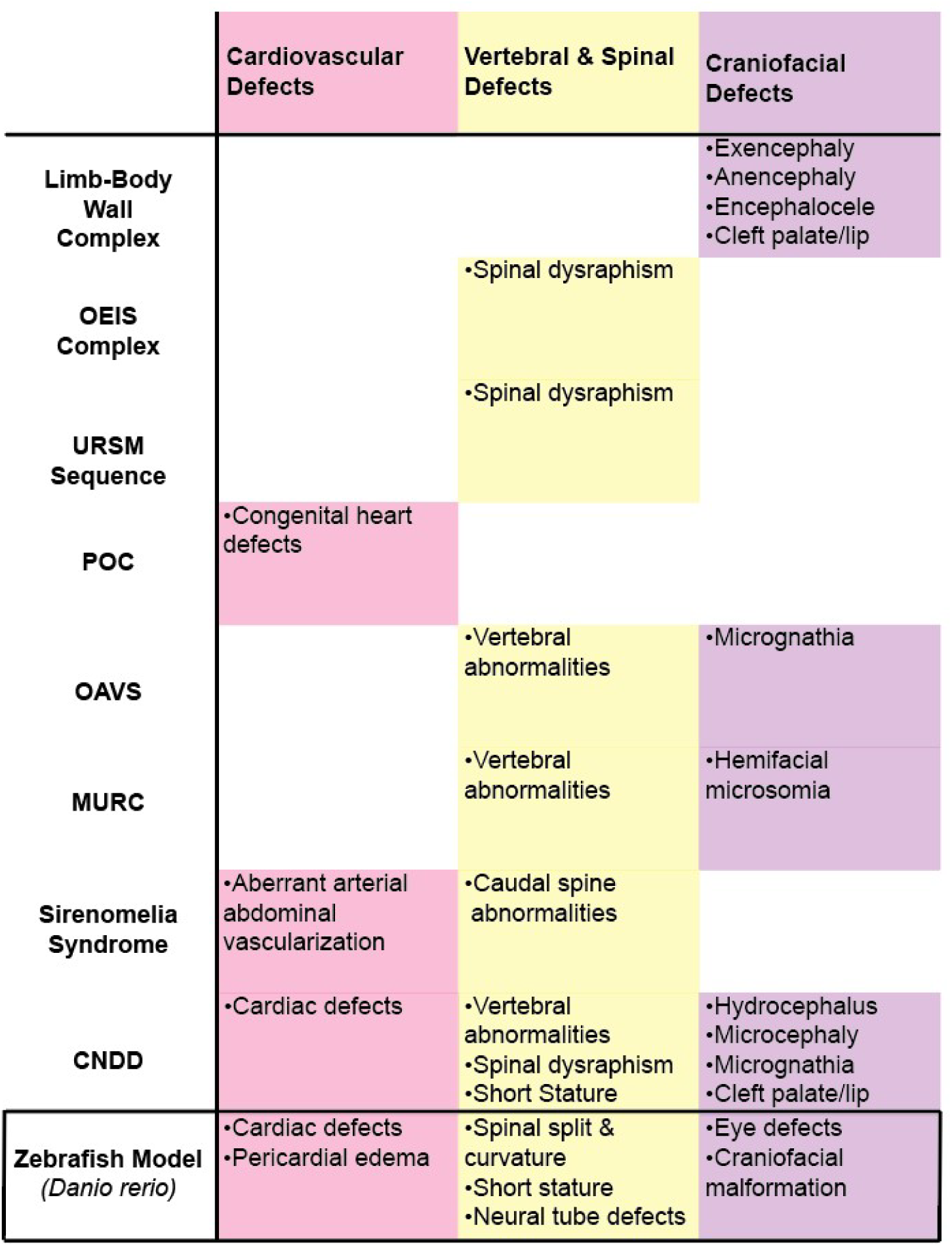
Congenital Defects and Zebrafish Phenotypes.

|  | Cardiovascular Defects | Vertebral & Spinal Defects | Craniofacial Defects |
| --- | --- | --- | --- |
| **Limb-Body Wall Complex** |  |  | •Exencephaly<br>•Anencephaly<br>•Encephalocele<br>•Cleft palate/lip |
| **OEIS Complex** |  | •Spinal dysraphism |  |
| **URSM Sequence** |  | •Spinal dysraphism |  |
| **POC** | •Congenital heart defects |  |  |
| **OAVS** |  | •Vertebral abnormalities | •Micrognathia |
| **MURC** |  | •Vertebral abnormalities | •Hemifacial microsomia |
| **Sirenomelia Syndrome** | •Aberrant arterial abdominal vascularization | •Caudal spine abnormalities |  |
| **CNDD** | •Cardiac defects | •Vertebral abnormalities<br>•Spinal dysraphism<br>•Short Stature | •Hydrocephalus<br>•Microcephaly<br>•Micrognathia<br>•Cleft palate/lip |
| **Zebrafish Model**<br>*(Danio rerio)* | •Cardiac defects<br>•Pericardial edema | •Spinal split & curvature<br>•Short stature<br>•Neural tube defects | •Eye defects<br>•Craniofacial malformation |

To assess whether ATDA impacts on NAD(H) levels in zebrafish, we measured NAD(H) levels in embryos treated with vehicle 2.5 mM ATDA, a dose similar to that used in mammals (Stewart et al., 1986). We found reduced NAD(H) levels in ATDA-treated embryos, indicating that ATDA impacts zebrafish NAD(H) metabolism (Fig. 2A). We also found that fewer ATDA-treated embryos at this developmental time point had hatched compared to vehicle-treated embryos (Fig. 2B), raising the possibility that ATDA induced defects in development. To determine whether zebrafish can model CNDD phenotypes in response to chemical-mediated NAD(H) disruption, we treated zebrafish embryos from four hpf with vehicle or a range of ATDA doses, and then assessed viability and

**Figure 2.**
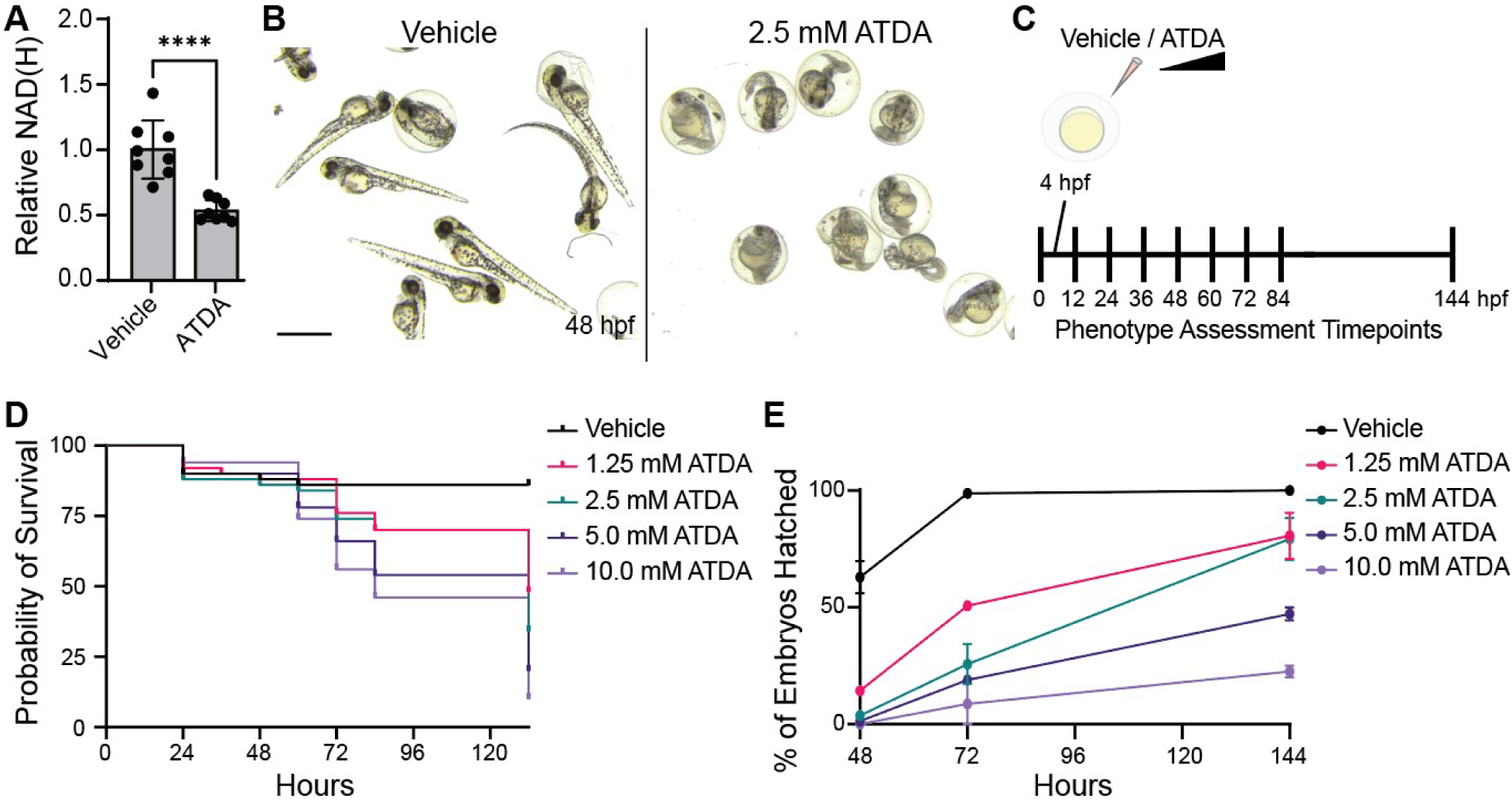
ATDA reduces zebrafish embryo survival and hatching success. A. NAD(H) was quantified in zebrafish embryos treated from 4-48 hpf with vehicle or ATDA. Error bars represent mean +/-SD of eight zebrafish in each condition (****=p<0.0001). B. Representative images of vehicle- and ATDA-treated embryos at 48 hpf. Scale bar = 1.0 mm. C. Schematic illustrating treatment and phenotype assessment protocol. Zebrafish embryos (4 hpf) were treated with vehicle or ATDA for up to 144 hours. D. Surviving embryos were quantified every 12 hours from 24-84 hpf and at 132 hpf. Data represent 50 embryos for each condition and are representative of two independent experiments. E. Hatched surviving embryos were quantified at 48, 72, and 144 hpf. Error bars represent mean +/- SD of two independent experiments, each containing 50 embryos for each condition.

CNDD-related phenotypes through 144 hpf (Fig. 2C). Over the time period of treatment, ATDA reduced zebrafish embryo survival and hatching success rates in dose-responsive manners (Fig. 2D,E). This is similar to the increase in fetal reabsorption observed in pregnant rats in response to NAD(H) inhibitors (Beaudoin, 1974, 1976) and the potential increase in spontaneous abortion rates in CNDD (Li et al., 2024), suggesting that loss of NAD(H) impacts zebrafish in a manner analogous to mammals.

### ATDA induces cardiac defects and tail malformations in zebrafish embryos

We also observed multiple additional phenotypes associated with CNDD and rodent models of NAD(H) disruption. Defects in tissues of mesodermal origin in CNDD and NAD(H) deficient rodent models include cardiac and vertebral defects (Table 2), which we also observed in our zebrafish model. We categorized pericardial edema according to severity (Fig. 3A, Table 1). The 72 hpf time point was chosen for analysis because the zebrafish larvae clearly displayed each phenotype of interest at this developmental stage (Boezio et al., 2023; Houk & Yelon, 2016). ATDA treatment induced a pronounced dose-dependent increase in pericardial edema (Fig. 3B), consistent with ATDA treatment impacting on cardiac development. We reasoned that CNDD and NAD(H) deficiency-related vertebral segmentation anomalies and defects such as caudal dysgenesis, hemivertebra, and block vertebra—each of which can cause abnormal spinal curvature—may manifest in zebrafish embryos as gradations of tail malformations. Indeed, we also observed a dose-dependent increase in tail malformation severity (Fig. 3C). Together, the developmental vertebral and cardiac defects observed in our experiments suggest that ATDA treatment of zebrafish embryos may accurately model CNDD and mirror NAD(H) deficient mammalian models.

**Figure 3.**
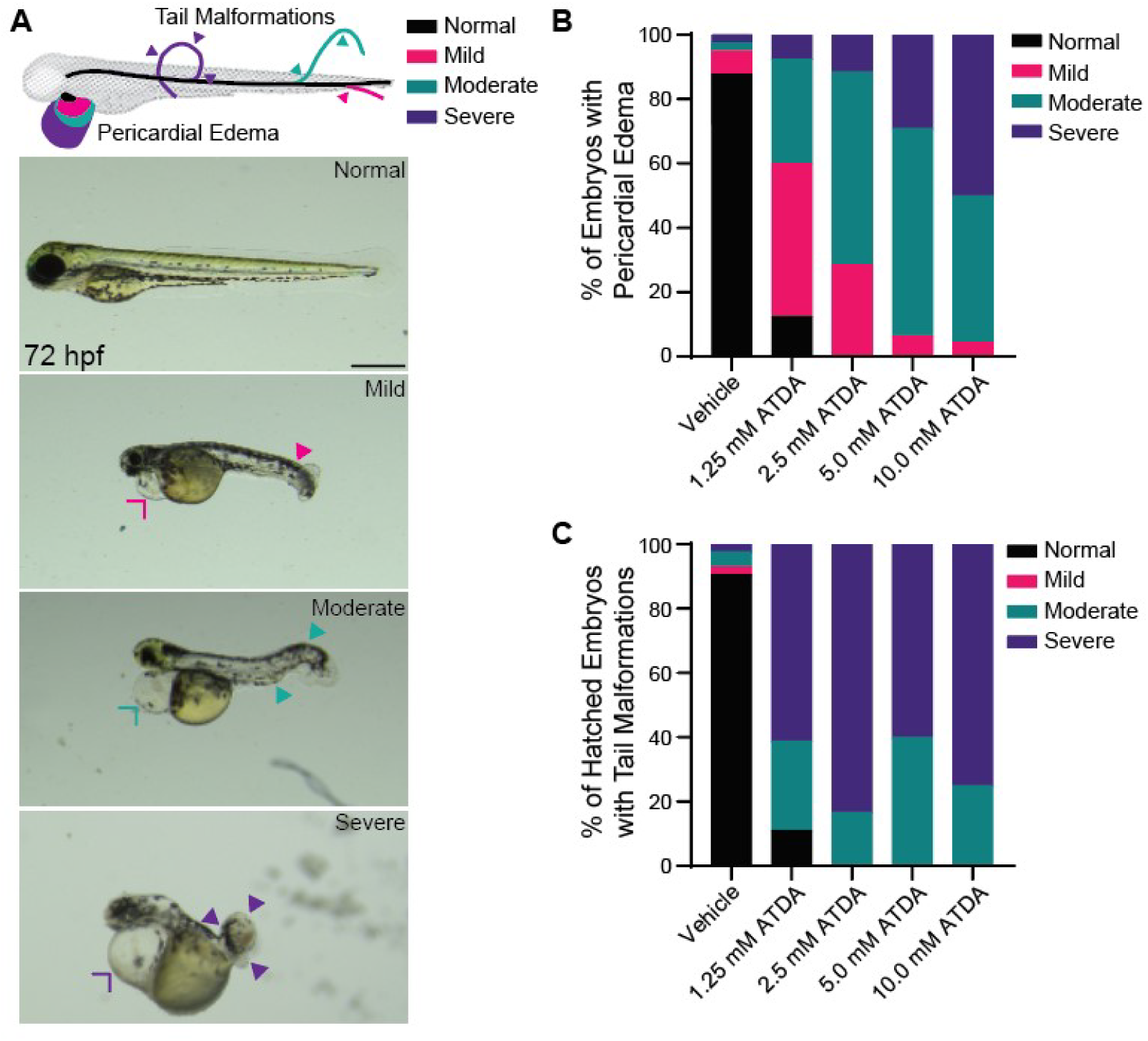
ATDA induces CNDD-like mesodermal abnormalities. A. Schematic representation and representative images of cardiac and tail abnormality scoring as described in Table 1. Closed arrowheads indicate tail curvatures. Open arrowheads indicate pericardial edema. Scale bar = 0.5 mm. B, C. 4 hpf zebrafish embryos treated with the indicated ATDA concentrations were assessed for pericardial edema (B) and tail malformations (C) at 72 hours. Data are from 50 embryos in each condition and are representative of two independent experiments.

### ATDA induces craniofacial abnormalities and spinal cord defects in zebrafish

CNDD and NAD(H) inhibition are additionally associated with ectodermal defects such as craniofacial abnormalities and neural tube defects (P. Mark & Dunwoodie, 2023; P. R. Mark, 2022). These broadly include facial findings such as hemifacial microsomia, hypertelorism, and micrognathia, along with skull and vertebral closure anomalies including anencephaly, exencephaly, and spinal dysraphisms, along with brain growth abnormalities including microcephaly and hydrocephalus (P. R. Mark, 2022) (Table 1). ATDA treatment induced head defects in zebrafish in a dose-dependent manner (Fig. 4A). We found that the developing zebrafish spinal cord parenchyma, which consists of proliferating neuroblasts and glioblasts, was disorganized and had reduced cellularity compared to control embryos (Fig. 4B). Finally, zebrafish embryos surviving to 144 hpf were substantially shorter than their vehicle-treated counterparts (Fig. 4C), reminiscent of the short stature common in CNDD (P. Mark & Dunwoodie, 2023).

**Figure 4.**
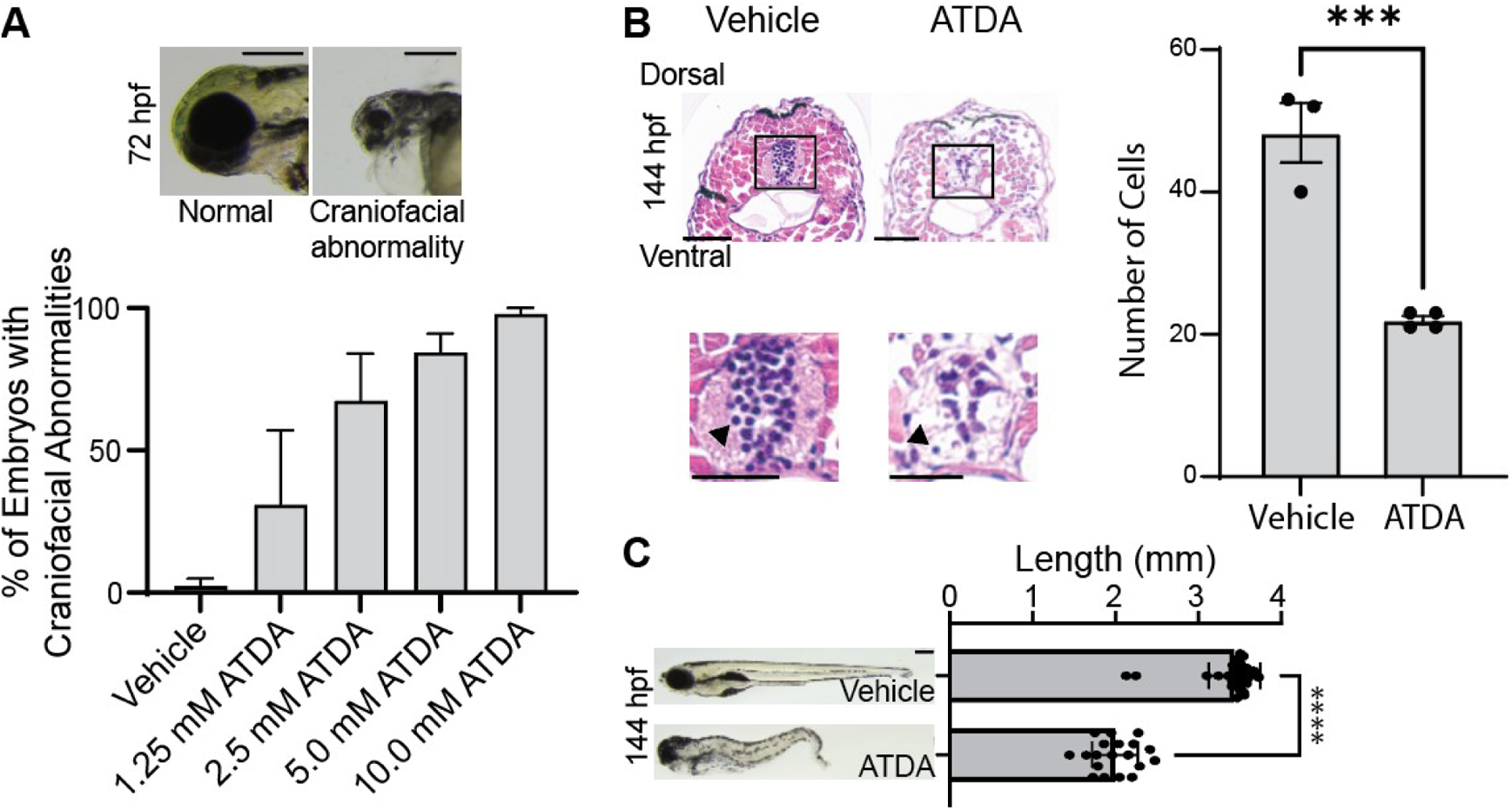
ATDA induces CNDD-like head defects and a failure of neural tube closure. A. Top: representative images of 72 hpf embryos with normal heads and with craniofacial abnormalities. Scale bar = 0.25 mm. Bottom: craniofacial abnormalities were quantified in embryos treated with the indicated ATDA concentrations at 72 hours. Data are means +/- SD of two independent experiments, each containing 50 embryos for each condition. B. Left: H&E-stained transverse sections at matched axial positions from embryos treated with vehicle or 2.5 mM ATDA for 96 hours. Scale bars = 50 µm. The spinal cord is boxed and an enlarged below. Arrowheads on enlarged spinal cord images indicate examples of basophilic cells that were quantified. Scale bars = 25 µm. Right: quantification of basophilic neuroblasts and glioblasts surrounding the spinal cord. Error bars represent mean +/- SD. Each point represents cells quantified cells from an embryo, and data are representative of two independent experiments (***=p<001). C. Length of embryos treated with vehicle or 2.5 mM ATDA for six days were measured using ImageJ tracing the body midline to account for body and tail malformations. Scale bar = 0.25 mm. Data are averages of at least 20 embryos in each condition and are representative of two independent experiments. Error bars represent mean +/- SD (****=p<0.0001).

### NAM rescues ATDA-induced phenotypes

To assess whether ATDA treatment-induced phenotypes were specific to the loss of NAD(H), we conducted a rescue experiment with nicotinamide (NAM), the initial metabolite in the NAD(H) salvage pathway (Fig. 1). NAM rescued NAD(H) levels in ATDA-treated embryos and appeared to rescue additional ATDA-induced developmental anomalies (Fig. 5A,B).

**Figure 5.**
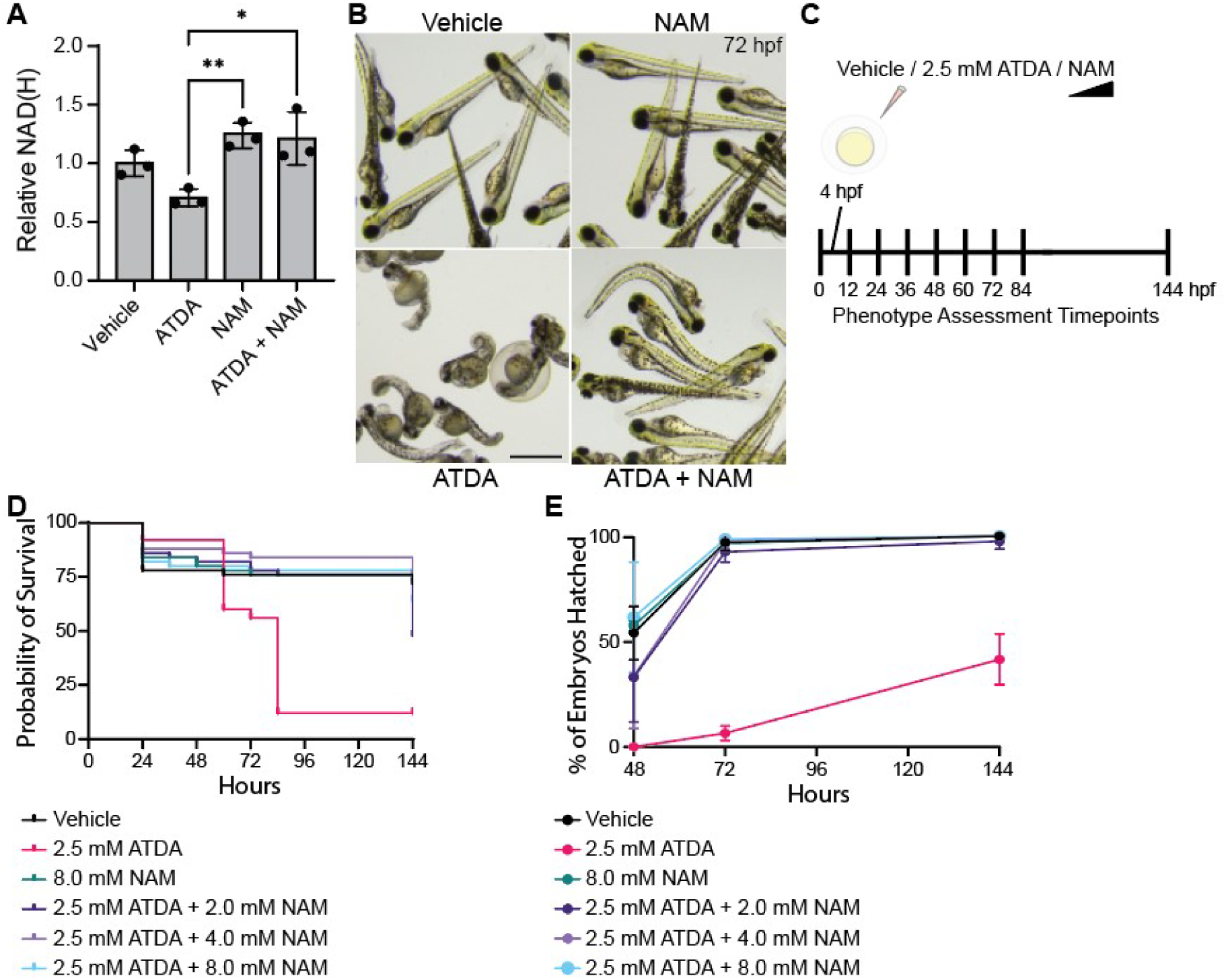
NAM administration rescues ATDA-induced impairment of zebrafish embryo survival and hatching success. A. NAD(H) was quantified in treated zebrafish embryos at 48 hpf. Error bars represent mean +/- SD of three zebrafish in each condition and are representative of two independent experiments (**=p<0.01; *=p<0.05). B. Representative images of treated embryos at 72 hpf. Scale bar = 1.0 mm. C. Schematic illustrating treatment and phenotype assessment protocol. Zebrafish embryos (4 hpf) were treated with vehicle, ATDA, NAM, or ATDA + NAM for up to 144 hours. D. Surviving embryos were quantified every 12 hours from 24-84 hpf and at 144 hpf. Data represent 50 embryos for each condition and are representative of two independent experiments. E. Hatched surviving embryos were quantified at 48, 72, and 144 hpf. Error bars represent mean +/- SD of two independent experiments, each containing 50 embryos for each condition.

To further assess of the impact of NAM on ATDA-mediated developmental defects, we co-treated 4 hpf zebrafish embryos with ATDA and a range of NAM concentrations to determine whether zebrafish also recapitulate mammalian models of NAD(H) disruption and rescue through metabolic intermediates (Beaudoin, 1976) (Fig. 5C). NAM restored embryo survival and hatching rates comparable to the vehicle (Fig. 5D,E). NAM co-treatment also rescued ATDA-induced pericardial edema and tail malformations (Fig. 6A-C) and craniofacial abnormalities (Fig. 6D) in dose-responsive manners. Further, NAM co-treatment rescued ATDA-induced spinal cord parenchyma disorganization and reduced cellularity (Fig. 6E), as well as zebrafish length (Fig. 6F). Together, these NAM rescue results indicate that the defects caused by ATDA are at least partially due to NAD(H) deficiency and further establish the utility of zebrafish as a model for assessing defects associated with depleted NAD(H) levels.

**Figure 6.**
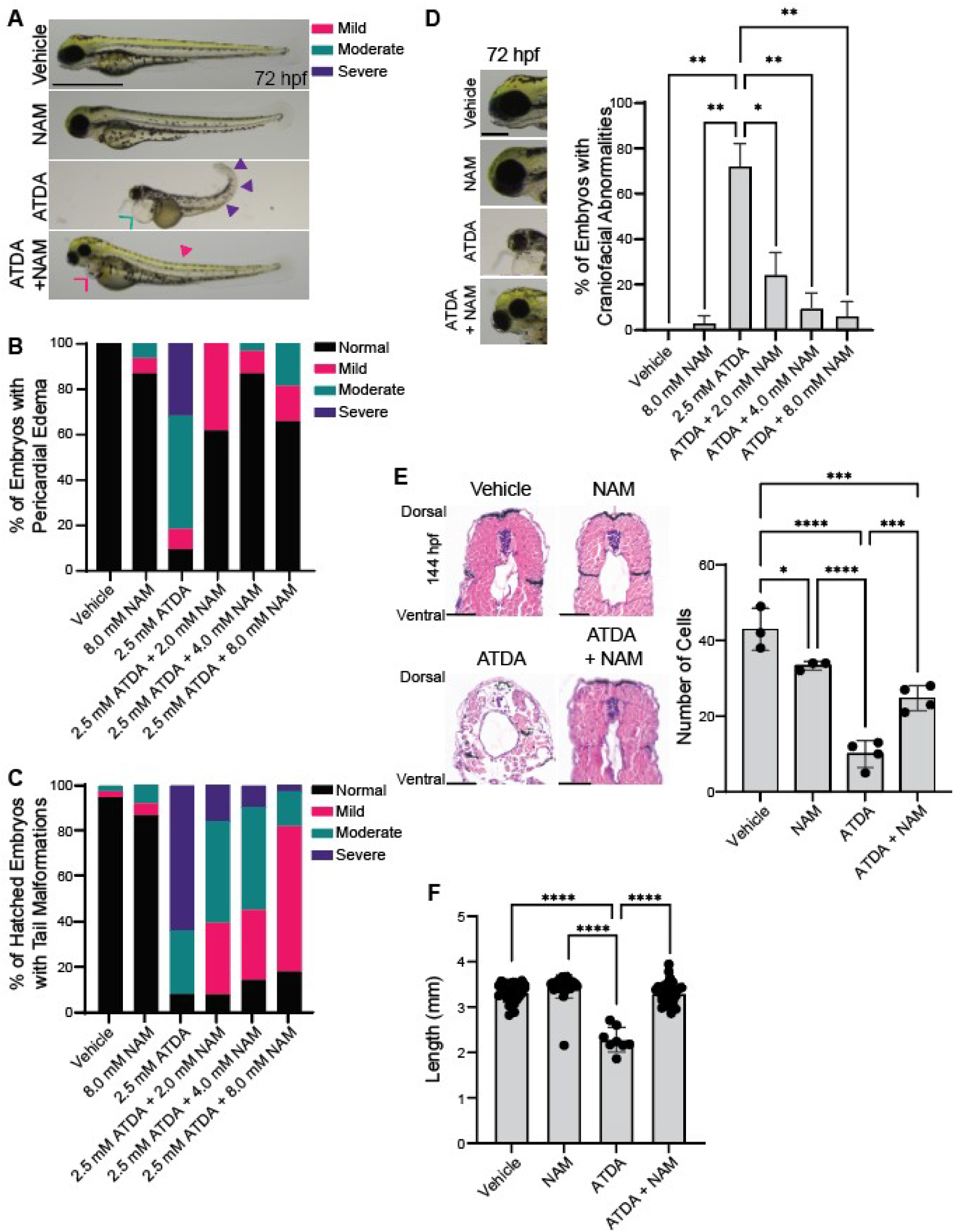
NAM administration rescues ATDA-mediated CNDD-like phenotypes. A. Representative images of phenotypes associated with vehicle, NAM, ATDA, or NAM + ATDA treatments. Closed arrowheads indicate tail curvatures. Open arrowheads indicate pericardial edema. Scale bar = 1.0 mm. B, C. 4 hpf zebrafish embryos treated with the indicated treatments were assessed for pericardial edema (B) and tail malformations (C) at 72 hours. Data represent 50 embryos in each condition and are representative of two independent experiments. D. Left: representative images of 72 hpf embryos with normal heads and with craniofacial abnormalities. Scale bar = 125 µm. Right: craniofacial abnormalities were quantified in embryos with 2.5 mM ATDA and the indicated NAM treatments at 72 hours. Data are means +/- SD of two independent experiments, each containing 50 embryos for each condition. E. Left: H&E-stained transverse sections at matched axial positions from treated embryos at 96 hours. Scale bars = 50 µm. Right: quantification of basophilic neuroblasts and glioblasts surrounding the spinal cord. Embryos were treated with vehicle, 8.0 mM NAM, 2.5 mM ATDA, or 8.0 NAM + 2.5 mM ATDA. Error bars represent mean +/- SD quantified cells from three embryos and are representative of two independent experiments (**** p<0.0001; ***p<0.001; **p<0.01; *p<0.05). F. Length of embryos treated with vehicle, 8.0 NAM, 2.5 mM ATDA, or 8.0 NAM + 2.5 mM ATDA for six days were measured using ImageJ tracing the body midline to account for body and tail malformations. Scale bar = 0.25 mm. Data are averages of at least 20 embryos in each condition and are representative of two independent experiments. Error bars represent mean +/- SD (****p<0.0001; ***p<0.001; *p<0.05).

## Discussion

CNDD has been linked to congenital anomalies and miscarriage in humans with biallelic variants in the NAD^+^ synthesis pathway genes *KYNU*, *HAAO*, and *NADSYN1* (P. Mark & Dunwoodie, 2023; Shi et al., 2017; Szot et al., 2020). Similar phenotypes are observed in mice with NAD^+^ synthetic precursor dietary deficiencies, and these effects are observed in mice both with or without NAD^+^ synthesis pathway genes variants (Cuny et al., 2020; Shi et al., 2017). This raises the possibility that low NAD^+^ levels may contribute to additional congenital anomalies. Many recurrent multiple malformation conditions exist which overlap in phenotype for which a genetic cause has not been identified, including limb-body wall complex (LBWC), pentalogy of Cantrell (POC), omphalocele-exstrophy-imperforate anus-spinal defects (OEIS) complex, oculoauriculovertebral spectrum (OAVS), Mullerian duct aplasia-renal anomalies-cervicothoracic somite dysplasia (MURCS), sirenomelia, and urorectal septum malformation (URSM) sequence, which all include VACTERL anomalies in their cardinal features (Adam et al., 2020; P. R. Mark, 2022). It has been hypothesized these conditions could be caused by a decreased maternal contribution of NAD^+^ through the yolk sac (P. R. Mark, 2022; Shi et al., 2017). Therefore, additional research assessing NAD^+^ restriction on development is necessary.

Thiadiazole compounds are used as fungicides, pesticides, antidiabetic, antimicrobial, and diuretic treatments, and are being studied as antiviral and anticancer agents (Babalola et al., 2024). Historically, studies using the thiadiazols, aminothiadiazole (ATD) and 2-Amino-1,3,4-thiadiazole (ATDA), in pregnant Wistar rats showed increased rates of embryonic lethality along with multiple birth defects in the offspring including neural tube, cardiac, limb, tail, kidney and palate anomalies (Beaudoin, 1974, 1976; Scott et al., 1973). These studies further revealed a connection between these teratogens and NAD^+^ metabolism by demonstrating that administration of NAM and metabolites from both NAD^+^ biosynthetic pathways could rescue the ATDA-induced defects in a dose and timing dependent fashion. The lethality and defects resulting from ATD and ATDA administration resemble those of CNDD and VACTERL patients. Therefore, NAD^+^ synthesis-disrupting teratogens such as ATDA remain viable tools to study NAD^+^-associated congenital defects. However, rodent studies are expensive and time consuming, and do not allow real-time observation of development. Here, we utilize zebrafish to model NAD^+^ deficiency during development to and recapitulate many of the systemic anomalies observed in CNDD and VACTERL patients. Our study demonstrates that zebrafish are a plausible model for studying the effects of NAD^+^ deficiency in growth and development.

Due to inherent differences between humans and zebrafish, phenotypic typing zebrafish for all anomalies found in humans with CNDD is not possible. However, humans and zebrafish share multiple overlapping organ system affected in CNDD and which are readily identifiable in our zebrafish model of NAD^+^ deficiency induced by ATDA (Fig. 3, Fig. 4A,C). These include craniofacial abnormalities, cardiac developmental defects, and tail defects which correlate with vertebral anomalies in humans (Table 2). Along with CNDD, these findings can be seen in the overlapping conditions LBWC, OAVS, POC, OEIS, URSM, and VACTERL association (Table 2). These conditions are defined by the non-random combination of these developmental abnormalities, and we have reliably produced these abnormalities in combination in our zebrafish model. We have additionally rescued these abnormalities by co-administering NAM with ATDA (Figs. 5, 6). Together, our findings demonstrate the utility of zebrafish as a model of these congenital anomalies and also provide further evidence for the possible role of low NAD^+^ levels in congenital anomalies.

Of interest, disrupting NAD^+^ metabolism in mice and rats produces neural tube defects at a higher frequency than humans with genetically derived CNDD disorder (Beaudoin, 1976; Cuny et al., 2023). Our results now demonstrate that this is also true for zebrafish (Fig. 4B). This could be due to an inherent increased susceptibility to neural tube defects of the animal species themselves. However, the environmental impact of ATDA teratogenicity in zebrafish and rats or dietary NAD^+^ precursor restriction in mice could influence neural tube development in a mechanism different than inherent genetic NAD^+^ deficiency seen in humans. Many recurrent malformation conditions for which there is no known genetic cause, such as LBWC, OEIS, and URSM, demonstrate frequent neural tube defects (Bergman et al., 2023; Mallmann et al., 2017; Nayak et al., 2023). Additionally, the environmental condition Diabetes Mellitus is a known risk factor for neural tube defects, has been linked to most of the recurrent malformation conditions listed above, and has been shown to impact NAD^+^ metabolism in mouse models (Castori, 2013; Chen, 2008; Salbaum et al., 2023).

In our zebrafish model, these findings, along with growth restriction, were all corrected by administration of the NAD^+^ precursor NAM in a dose-dependent fashion, demonstrating the importance of NAD^+^ in the growth and development of these organ systems. Future studies in zebrafish assessing additional CNDD- and VACTERL-impacted organs and tissues such as kidney and limbs may also contribute to our understanding of these congenital anomalies. Because cellular and molecular markers of development are well-described in zebrafish, future cellular and molecular investigations in zebrafish may also shed light on the mechanisms driving these anomalies.

## Conclusions

Here, we have generated a zebrafish model of NAD^+^ synthesis inhibition to model congenital abnormalities arising from suppressed developmental NAD^+^. ATDA treatment induced cardiac, craniofacial, tail, body length, and neural tube defects. Each of these defects was reversed by administration of NAM, a metabolite in the NAD^+^ salvage pathway. Our work described here demonstrates the utility of zebrafish as a model to further investigate the ontogeny of CNDD and other congenital anomalies that may result from restricted NAD^+^ during development.

## Author Contributions

**Visakuo Tsurho**, **Carla Gilliland**, and **Jessica Ensing**: Methodology, Investigation, Writing - Review & Editing. **Elizabeth Vansickel**: Methodology, Writing - Review & Editing. **Nathan J. Lanning**: Methodology, Investigation, Formal Analysis, Writing – Original Draft, Writing - Review & Editing, Visualization, Project Administration. **Paul R. Mark**: Conceptualization, Methodology, Supervision, Project Administration, Writing – Original Draft, Writing - Review & Editing, Funding Acquisition. **Stephanie Grainger**: Conceptualization, Methodology, Supervision, Project Administration, Writing – Original Draft, Writing - Review & Editing, Funding Acquisition.

## Acknowledgements

Research reported in this publication was supported by the National Institute of General Medical Science under Award Number R35GM142779 (SG). The content is solely the responsibility of the authors and does not necessarily represent the official views of the National Institutes of Health. C. Bradfield is thanked for zebrafish husbandry.

## Data and materials availability

All materials are available upon reasonable request.

